# A conditional, myeloid-cell specific estrogen receptor α deletion reprograms the liver immune microenvironment and impedes the growth of colon carcinoma liver metastases

**DOI:** 10.64898/2026.08.28.747896

**Authors:** Orçun Haçariz, Megan Kalaw, Qin Yang, Stephanie Perrino, Pnina Brodt

## Abstract

Liver metastases (LM) remain a major cause of death from different cancer types, in particular malignancies of the gastrointestinal tract. Liver metastases predict a poor response to immunotherapy due, among others, to the immunotolerant microenvironment (ME) of the liver and loss of local and systemic cytotoxic T cells. Thus, strategies that can reprogram the immune ME of the liver and restore cytotoxic T cell reactivity are being sought. We previously reported that estrogen signaling blockade impedes the growth of LM by reducing MDSC accumulation and monocyte/macrophage polarization. The aim of this study was to elucidate the underlying mechanism(s) and assess whether estrogen signaling in the myeloid lineage was driving the immunotolerant ME of LM. To this end, we generated mice with conditional myeloid cell-specific deletions of estrogen receptors (ER)α or ERβ and analyzed in these mice the effect of ER loss on the liver immune ME and the outgrowth of LM. In mice with ERα, but not with ERβ deletion, we observed a marked reduction in the growth of murine colon carcinoma MC-38 liver metastases as compared to their respective controls. Flow cytometry and immunohistochemistry revealed a decrease in macrophages that were polarized to the pro-tumorigenic M2-like phenotype and a concomitant increase in activated CD8^+^ T and NK cells relative to controls. Bulk RNAseq analysis performed on hepatic immune cells infiltrating the liver revealed changes in the expression of key cytokines/chemokines mediating immune cell recruitment, activation and polarization, including *Ccl5* (upregulated) and *Csf1* (downregulated). Taken together, the data suggest that ERα signaling in myeloid-derived cells programs the immune landscape and contributes to an immunosuppressive and metastases-growth permissive ME in the liver.

## Introduction

Liver metastases, cancerous growths that have spread to the liver from other sites, remain a major cause of death from different cancer types, particularly malignancies of the gastrointestinal (GI) tract (1). Recently, the incidence of GI cancers, in particular colorectal cancer, has been increasing in younger adults (< 50 yr of age) (2) but the factors responsible for this trend are still under investigation (3). Recent studies suggest that increased obesity, diabetes and chronic inflammation in the young population, as well as the microbiota may contribute to this trend (3, 4), however the role of sex hormones has not to date been investigated.

Immunotherapy (IT) has become a standard of care (SOC) for different malignancies, including melanoma and small cell lung cancer, where therapeutic efficacy has been demonstrated. However, the therapeutic benefit from IT is diminished when liver metastases are present (5, 6). This lack of response to immunotherapy was attributed, at least in part, to T cell suppression by other immune cells that infiltrate the TME and release immunoregulatory cytokines such as IL-6 and/or IL-10 (7) or to the elimination of activated T cells by F4/80+ CD11b+CCR2+ macrophages with an M2-like signature via Fas/FasL-induced apoptosis (8).

Bone marrow derived cells of the myeloid lineage, including neutrophils, macrophages, dendritic cells and myeloid derived suppressor cells (MDSCs) are recruited into the tumor microenvironment (TME) of LM. The accumulation of monocytic (M-MDSCs) and granulocytic (G-MDSCs) MDSC results in potent immunosuppressive activity (1, 9). MDSCs strongly suppress adaptive T-cell mediated immunity by releasing immunosuppressive factors such as TGFβ and arginase (10). Monocyte/macrophages recruited to the liver can polarize to a pro-tumorigenic phenotype and release tumor promoting factors such as VEGF and MMP-9 and T cell suppressing factors such as IL-10, TGFβ and arginase. Together, MDSCs and M2-like macrophages in the TME are currently thought to be major barriers to successful cancer immunotherapy (11).

Estrogen is a pleiotropic steroid hormone that transmits its signals to regulate gene transcription via its nuclear receptors ERα and ERβ (12). Although the regulation of MDSCs by estrogen has been documented in several studies (13), the underlying mechanism(s) including the relative roles of the two receptors in the context of liver metastasis is not clear.

Our previous studies revealed that estrogen plays a key role in regulating a pro-metastatic immune microenvironment in the liver (14, 15). We reported that liver metastasis of colon, and lung carcinoma cells was inhibited in TNFR2 null female, but not male mice (16), and this was linked to estrogen signaling (14), suggesting that estrogen may regulate TNFR2 expression and signaling in immune cells (14), as was also reported for other cell types (17, 18). In ovariectomized mice, a marked reduction was observed in colorectal, lung and pancreatic carcinoma liver metastases that was associated with reduced liver MDSC accumulation, increased IFN-γ and Granzyme B production by CD8^+^ T cells and reduced TNFR2, IDO2, TDO and Serpin B9 expression levels in the liver (14). Subsequently we reported that estrogen played a role in macrophage polarization to the M2 phenotype and inhibition of estrogen signaling by Fulvestrant potentiated the tumor-inhibitory effect of immunotherapy on hormone-independent and immunotherapy-resistant metastatic cancer (15). It remains unclear however, whether estrogen acts directly on immunosuppressive cells recruited into the liver or these cells receive their cues from other resident and recruited cells in an estrogen-regulated fashion. To address this question, we generated mice with a myeloid-specific deletion of ERα or ERβ and explored the effect of these deletions on the immune ME of LM and on the outgrowth of liver metastases.

## Results

### Reduced liver metastases in mice with a myeloid specific ERα but not ERβ deletion

To assess whether the selective loss of ER expression in myeloid cells affected the establishment of colon cancer liver metastases, we treated the Cre-Lox mice (Cre+ and Cre- as controls) for 5 days with tamoxifen via oral gavage to activate Cre recombinase and 12-14 days after the last tamoxifen injection (to ensure no direct effect of tamoxifen) injected them with 2X10^5^ murine colon carcinoma MC-38 cells via the intrasplenic/portal route. Mice were euthanized and livers removed 17-23 days later and visible metastases on the surface of the livers enumerated. We found a marked reduction in the number of liver metastases in mice with a myeloid-specific ERα (ERα KO) (P < 0.05) but not ERβ (ERβ KO) deletion relative to their respective controls (**Figure 1 A, C and D**). The number of LM in ERα KO mice was also significantly lower than in ERβ KO mice (**Figure 1A right and left**). Intriguingly, we found in the latter group an overall increase in the number of liver metastases relative to the respective controls, although the difference didn’t reach statistical significance. The size of individual metastases was similar in all groups (**Figure 1B**), suggesting that the loss of ERα expression in myeloid cells caused an early elimination of invading tumor cells rather than a reduction in the expansion of individual micrometastases. Taken together, these results suggested that (i) ERα signaling in myeloid-derived cells played a tumor-promoting role in the liver ME and (ii) ERα and ERβ played distinct and possibly opposing roles in the immune ME of LM. They also indicated that tamoxifen administered orally several weeks prior to tumor cells injection did not in itself result in a reduction in the number of liver metastases.

**Figure 1.**
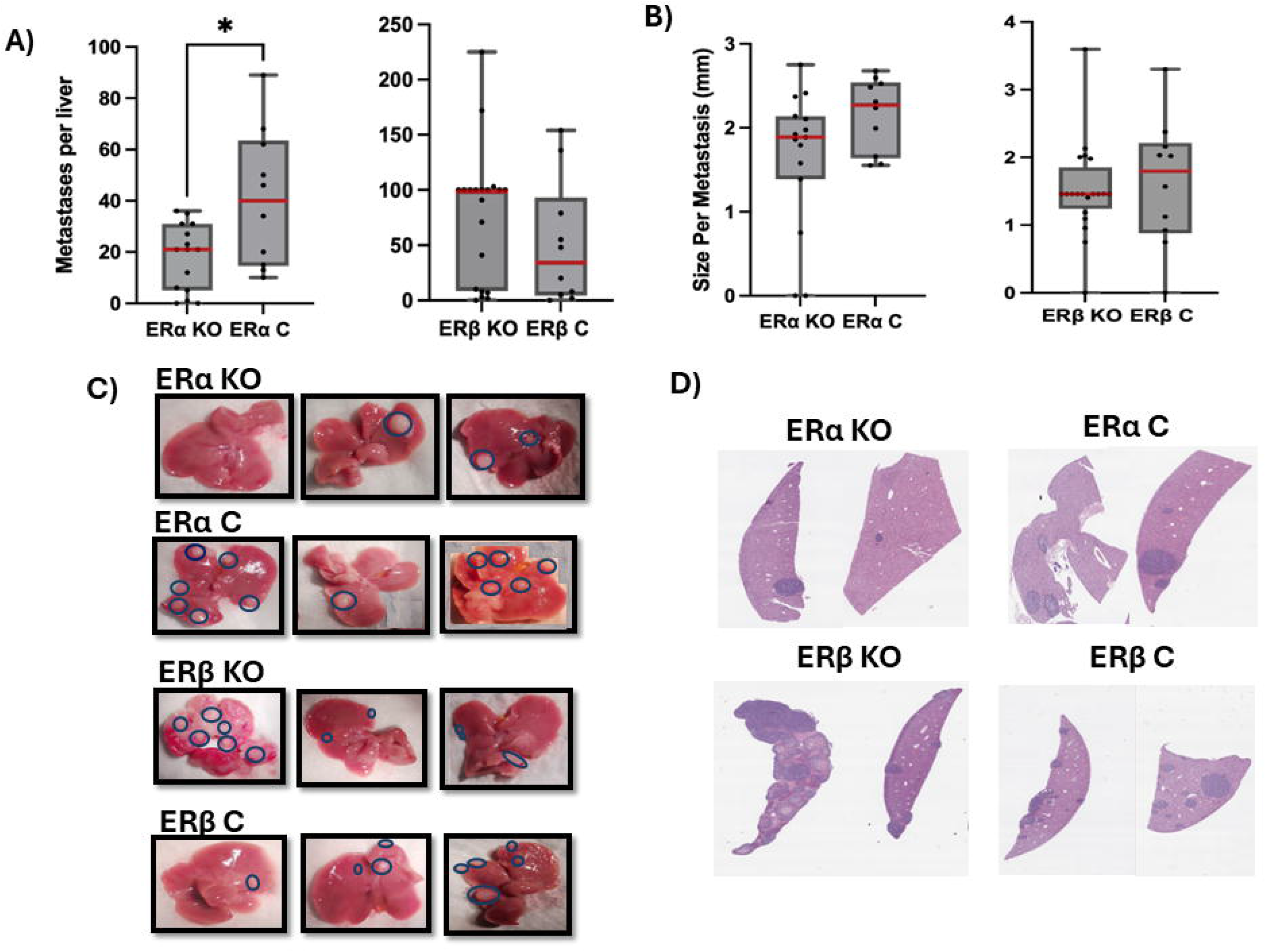
Reduced colon cancer liver metastasis in mice with a myeloid cell - specific deletion of ERα. The number of metastases per liver and median are shown in **(A)** and the sizes per visible metastasis (in mm) are shown in **(B)**. Representative livers are shown in **(C)**, H&E- stained formalin-fixed and paraffin-embedded sections for the ERα KO, ERβ KO and respective control groups are shown in **(D)**. The number of metastases per liver (but not their sizes) was significantly lower in the ERα KO group as compared to the Cre^-^ littermate controls (ERα C) (P < 0.05). The median number and mean metastases size in ERβ KO mice were not significantly different from those in the Cre- controls. The results are pooled data from 2 or-3 separate experiments. Total number of mice per group was: ERα KO; n=15, ERα C; n = 10, ERβ KO; n=20 and ERβ C; n=10.

### Reduced ER**α** expression in bone-marrow derived myeloid cells increases M1 polarization in liver-recruited macrophages

We previously reported that estrogen depletion by ovariectomy altered the macrophage population in the liver TME, significantly increasing the M1:M2 macrophage ratio relative to controls in both resident and recruited macrophages (15). Having mice with a selective depletion of ERα in bone-marrow derived myeloid cells provided an opportunity to assess whether ERα signaling in the liver-recruited monocyte/macrophages was directly responsible for the altered phenotype. Macrophages isolated from the liver 8-10 days post tumor cell injection were analyzed by flow cytometry. We found no difference in the proportions of CD11b^+^CD68^+^CCR2^+^ recruited macrophages in ERα KO relative to control mice, where they constituted approximately 20% of the total viable cells (**Supplementary Figure 2A**). However, among these cells, the proportion of CD163^-^/CD38^+^ M1-like macrophages was significantly higher in livers derived from ERα KO mice than in controls (P < 0.05), while the proportion of the CD163^+^/CD38^-^ M2- like subset was not significantly altered, resulting in a significantly elevated M1:M2 macrophage ratio (**Figure 2A**). These results were confirmed when liver cryostat sections derived from the livers of tumor injected mice were immunostained, revealing an increase in the proportion of M1-like macrophages (**Figure 2B**). In contrast to these results, we found no change in the polarization status of macrophages recruited to the liver IME in mice with a myeloid specific ERβ deletion (**Figures 2A&B**). Taken together, these results suggested that the loss of ERα but not ERβ expression retained a larger proportion of the recruited macrophages in a proinflammatory M1-like polarization state.

**Figure 2.**
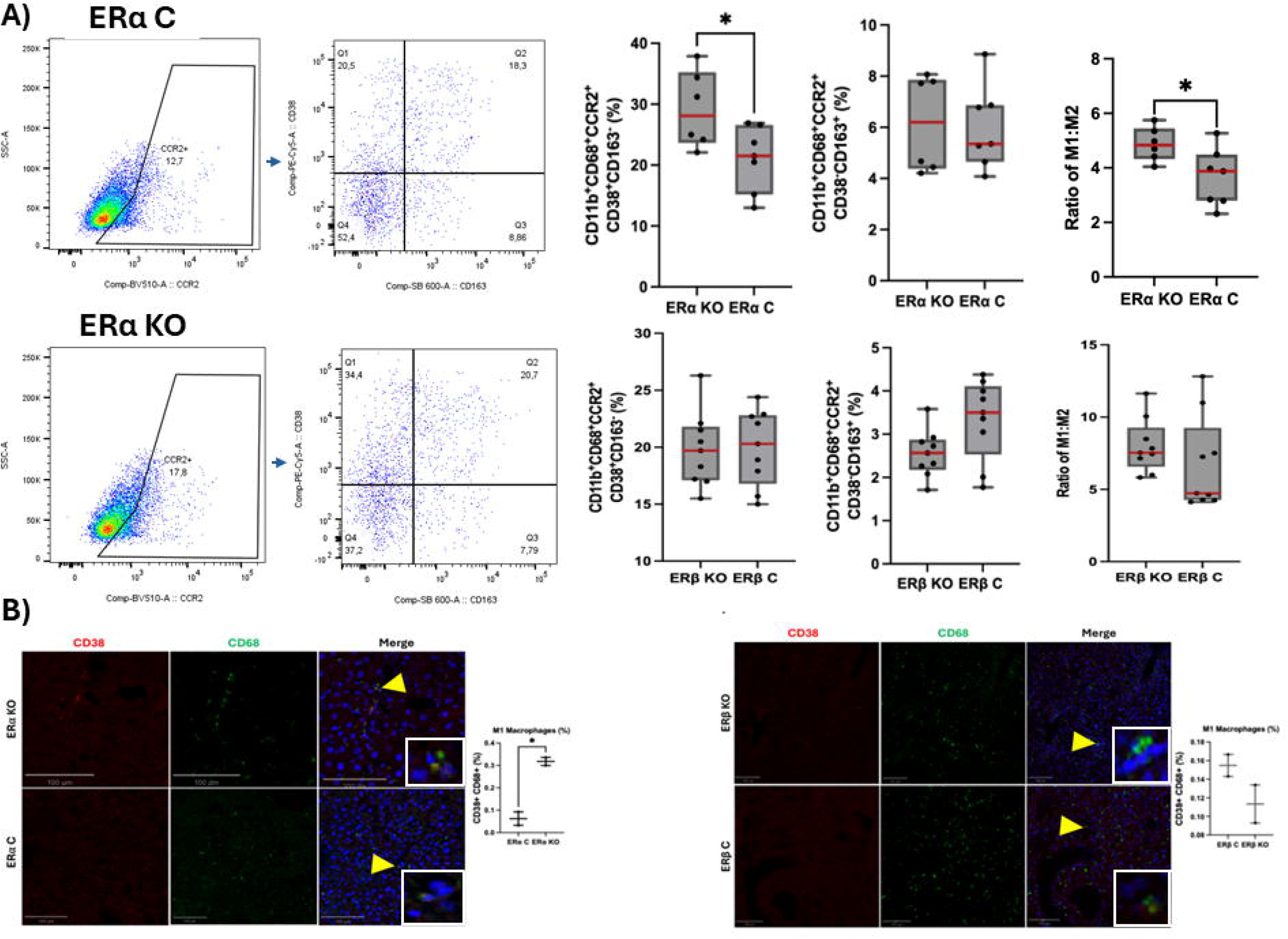
Increased M1:M2 macrophage ratio in the liver IME of mice with a myeloid cell- specific ERα (but not ERβ) deletion. Shown in **(A)** are results of flow cytometry analysis of liver recruited macrophages 8 days post intrasplenic/portal injection of MC-38 cells and the M1:M2 ratios (right). The M1:M2 ratio was significantly higher in ERα KO mice as compared to the controls based on 2 separate experiments and 3-5 livers per group in each experiment (P < 0.05). Shown in **(B)** are representative images of IF staining performed on cryostat sections derived from livers of MC-38 injected mice 8 or 9 days post injections. Yellow arrows denote M1-like macrophages. On the right of the images are results of the quantification of M1-like macrophages based on 15 fields counted in a total of 3 sections per group and expressed as % of all CD68+ macrophages per field.

### Loss of ER**α** expression in myeloid cells results in increased CD8^+^ T cell and NK cell activation in the liver TME

Having observed an increase in the M1:M2 macrophage ratio in the ERα KO mice, it was of interest to determine whether this impacted T and NK cell recruitment and/or activation states in the liver IME. Analysis of hepatic immune cells (HIC) by flow cytometry revealed no difference in the recruitment of CD3e^+^/CD8b^+^ T cells or the numbers of CD3e^-^/NKp46^+^ NK cells in the liver TME of these mice relative to controls (**Supplementary Figure 2B&C**). However, for both lymphoid populations, a significant increase was observed in the proportion of activated cells as determined by enhanced expression of both granzyme B (GZB) and interferon γ (IFNγ) in both cell types (**Figures 3 A&B**). We confirmed the enhanced cytotoxic activity of isolated T cells against cultured MC-38 cells using the Incucyte system. When CD8^+^ T cells isolated from the mice 8 days post tumor cell injection were incubated with the tumor cells, a significant increase in tumor cell kill was observed with T cells derived from ERα KO mice relative to controls (**Figure 3C**).

**Figure 3.**
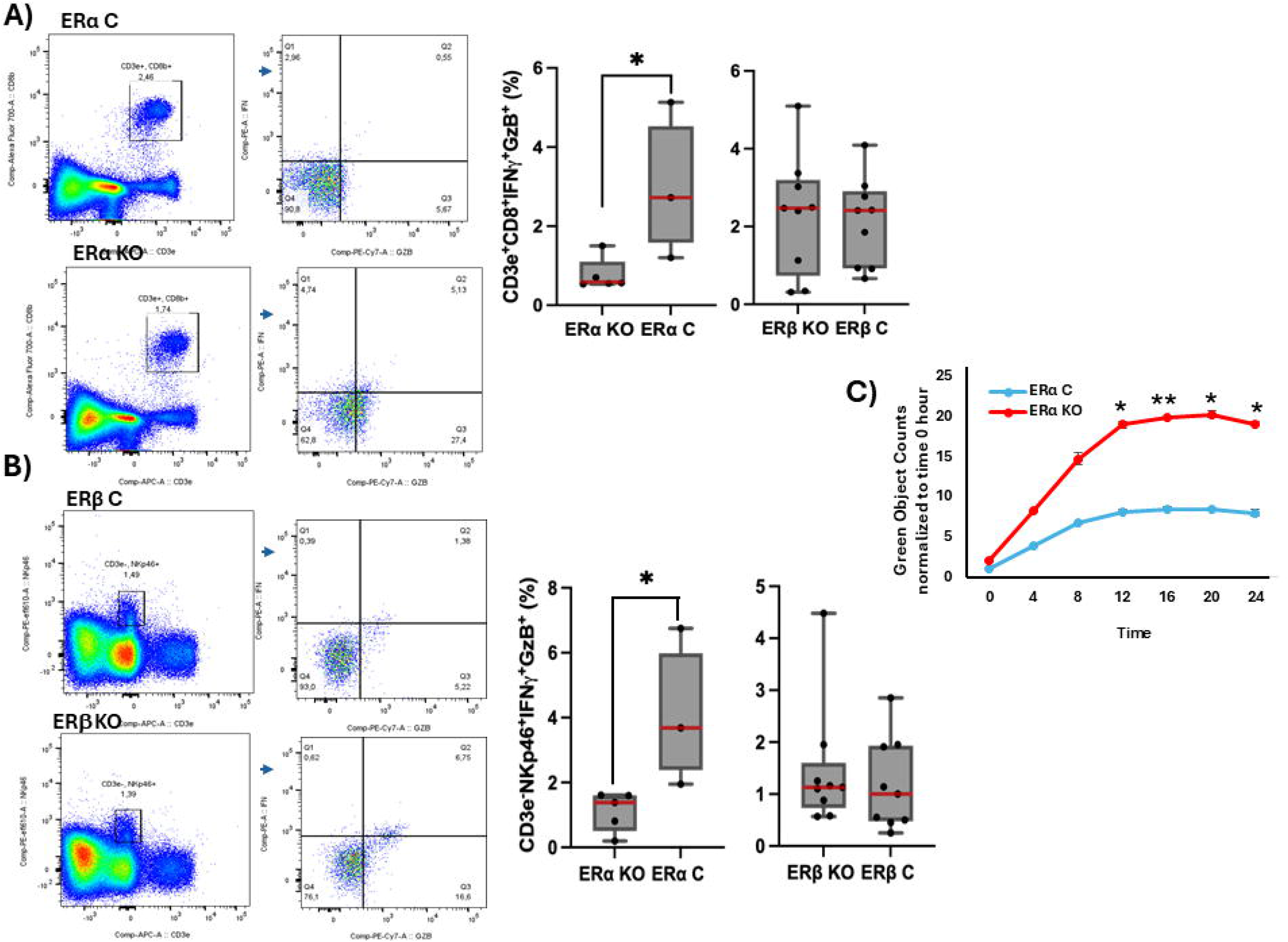
Increased T and NK cell activation in mice with myeloid cell specific ERαdeletion. Shown in **(A&B)** are results of flow cytometry analyses performed on immune cells isolated from the livers of mice 7 or 8 days post injection of 2 × 10^5^ MC-38 cells. Profiles of CD8^+^ T cells are shown in **(A)** and of NK cells in **(B)**. A significant increase in activated T and NK cells relative to respective controls was found in Erα, but not in ERβ KO mice (P < 0.05, based on 2 experiments and 3-5 mice per group). Results of cytotoxicity assay performed using the Incucyte live cell imaging system are shown in **(C)**. The results are normalized to time 0 (analysis based on triplicates and 2 mice per group). MC-38 cell death peaked at 12 hr and was significantly higher following incubation with CD8^+^ T cells derived from ERα KO mice as compared to controls. *p < 0.05, **p<0.01.

### Myeloid cell-specific deletion of ER**α** alters the transcriptomic landscape of the liver metastases-associated immune cells

Our results have clearly shown that loss of ERα expression in myeloid-derived cells altered the IME of liver micrometastases. To investigate transcriptomic changes in the IME underlying these changes, we employed a global approach and subjected the HIC to bulk RNA sequencing using 3 samples of total harvested HIC per group (ERα competent and KO mice). Approximately 40 million reads were obtained per sample. Principal component analysis (PCA) confirmed a clear segregation between the different experimental groups (**Figure 4A**). Differentially expressed gens in the groups are listed in **Supplementary Data 1**. A total of 1368 genes were found to be differentially expressed between IHC of ERα KO and control mice (FDR < 0.05), as depicted in the Volcano plot (**Figure 4B**). A significant reduction in *Esr1* expression in HIC derived from ERα KO mice as compared to the controls was confirmed by qPCR performed on the same RNA pool (**Figure 4C**).

**Figure 4.**
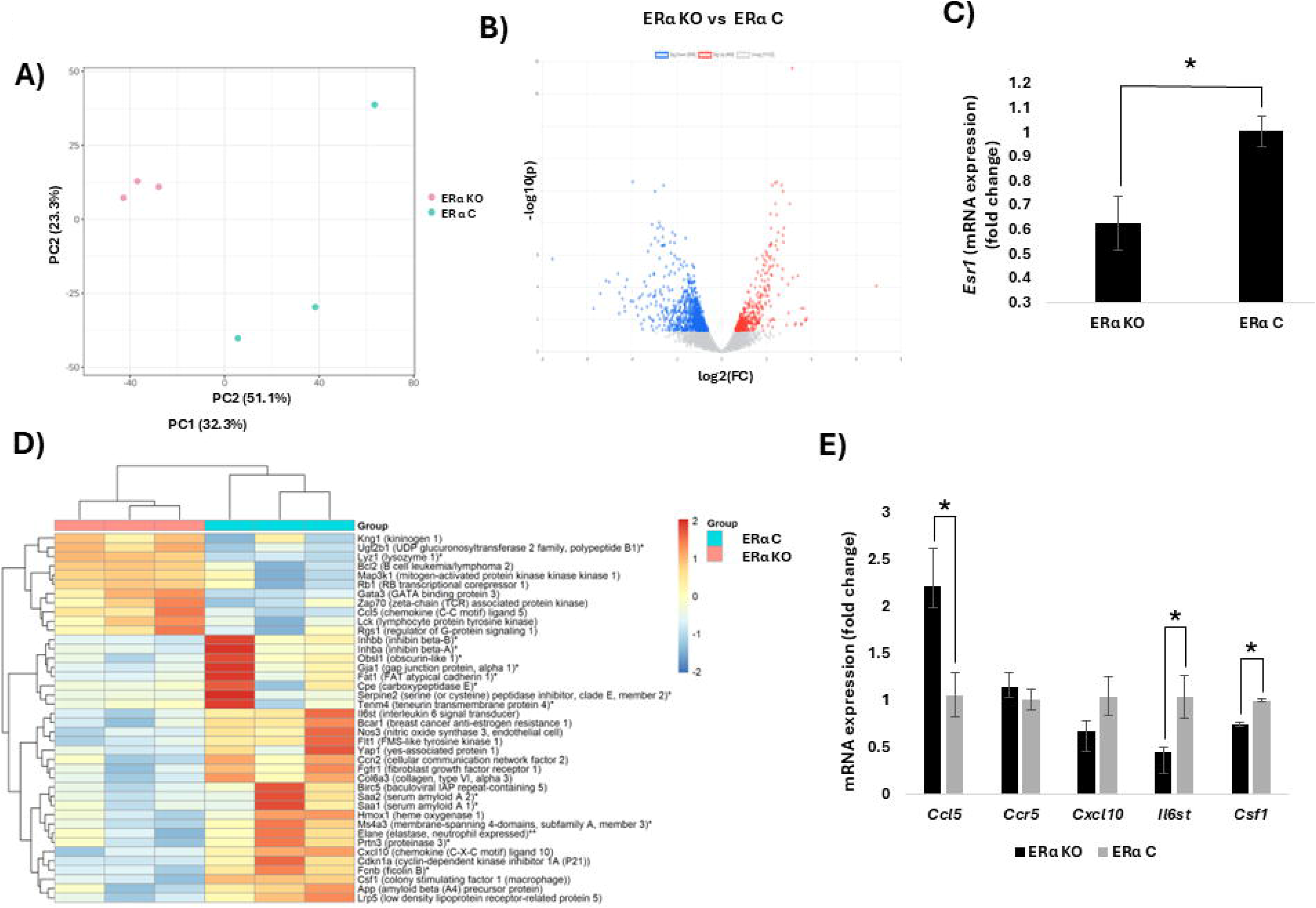
Altered transcriptomic landscape in the TME of mice with a myeloid cell specific ERα deletion. Shown in **(A)** is the PCA depicting inter and intra-group variations and a clear separation between the experimental groups. Shown in **(B)** is a Volcano plot with up and downregulated DEGs (FDR < 0.05) in the comparison between ERα KO and ERα C - derived HIC. Shown in **(C)** are qPCR results for estrogen receptor α (*Esr1*) expression levels in ERα KO and ERα C-derived HIC. *Esr1* expression levels were significantly lower in ERα KO in comparison with the controls (P < 0.05, n=3). In the Heatmap shown in **(D)** known transcriptional targets of *Esr1* (e.g. *Csf1*), highly regulated DEGs (FC=|3|, (FDR < 0.05), and estrogen-related genes (FDR < 0.05) identified by GeneCards are highlighted. *-denotes DEGs related to estrogen signaling as identified by GeneCards: **denote transcripts that are both highly regulated and identified by GeneCards as related to estrogen signaling. Genes with sum counts greater than 200 in each group were included. Results of qPCR analysis shown in **(E)** confirm differential expression of transcripts of interest (*Ccl5*, *Cxcl10*, *Il6st*) and transcripts of potential relevance (e.g. *Ccr5,* the CCL5 receptor). The results are expressed as means±SE based on 3 biological replicates per transcript. The expression of *Ccl5* was increased and of *Il6st* and *Csf1* decreased in ERα KO as compared to controls (P < 0.05). There was no statistical difference in the expression levels of *Cxcl10* or *Ccr5*.

Among the genes that were differentially expressed in HIC derived from ERα KO mice relative to the controls (FDR < 0.05) were known estrogen regulated transcripts (as determined by GeneCards or based on published reports), although the changes in expression levels did not always follow the predicted patterns (**Figure 4D**). They included chemokine (C-X-C motif) ligand 10 (*cxcl10*) and chemokine (C-C motif) ligand 5 (*ccl5*), whose protein products were implicated in the recruitment and activation of innate and adaptive immune response cells to the tumor ME (15). The differential expression of these genes was subsequently confirmed by qPCR (**Figure 4E**). While expression of *Ccl5* was significantly increased in HIC derived from ERα KO mice (in agreement with our previous findings in ovariectomized mice (15)), the expression of *Il6st,* the signal transducer of the immunosuppressive cytokine IL-6 and of colony stimulating factor 1 (*Csf1*) was decreased in ERα KO mice relative to controls, as also confirmed by qPCR (P < 0.05), consistent with an immunosuppressive role of ERα within the TIME of the liver.

Analysis by overexpression gene enrichment analysis (OA) of Gene Ontology-Biological Processes (GO-BPs) (**Supplementary data 2**) revealed that the differentially expressed genes (FDR < 0.05) in ERα KO mice were enriched mainly in those involved in cellular organization and the immune responses (**Supplementary Figure 3**). The majority of genes involved in positive regulation of the immune response (GO:0050778, FDR < 0.001), T cell activation (GO:0042110, FDR < 0.01) and interferon gamma production (GO:0032609, FDR < 0.01) were upregulated in ERα KO as compared to controls (**Supplementary Figures 4A-C**). The enriched biological pathways (KEGG) analysis (**Supplementary data 2**) revealed upregulation of transcripts involved in NK cell mediated cytotoxicity (FDR < 0.01) in these mice (**Supplementary Figure 4D**) and a similar pattern was observed for T-cell receptor activity (FDR < 0.05) (**Supplementary Figure 4E**).

Furthermore, gene set enrichment analysis (GSEA) (**Supplementary data 3**) revealed a positive regulation of immune cell activation while developmental processes were negatively regulated in ERα KO immune cells (**Figure 5A**). Of particular relevance, this analysis revealed a negative regulation of the macrophage colony stimulating factor biological process (M-CSF, CSF-1) (NES: -3.89, FDR = 0) in ERα KO immune cells **(Figure 5B)**, in line with documented evidence that *Csf1* is transcriptionally regulated by ERα (19), and a significant downregulation of *Csf1* in immune cells derived from ERα KO mice was also confirmed by qPCR (**Figure 4E**). On the other hand, the natural killer cell mediated cytotoxicity biological pathway was identified as positively regulated in these cells (NES: 2.65, FDR < 0.001) (**Supplementary data 3** and **Figure 5C**), in line with the results of the OA analysis **(Supplementary Figure 4D)** and our flow cytometry data (**Figure 3**).

**Figure 5.**
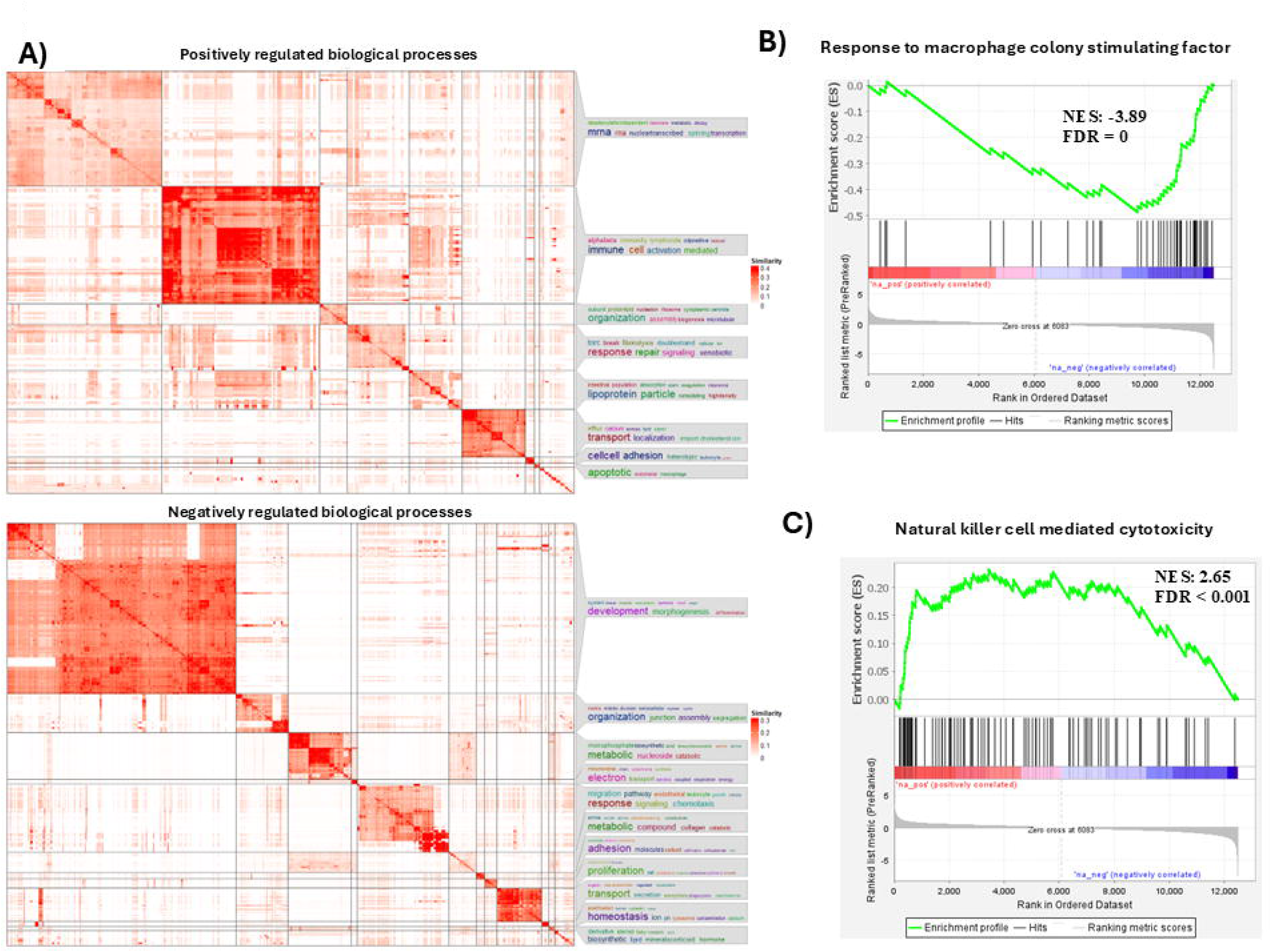
GSEA analysis identifies biological processes differentially regulated in the immune ME of liver metastases in mice with myeloid cell specific ERα deletion. Shown in **(A)** are positively (e.g. immune responses-related) and negatively (e.g. development-related) regulated biological events in ERα KO (compared to ERα C) based on the clustering of the enriched GO-BPs (P < 0.05) by simplifyEnrichment. In **(B)** are Enrichment plots demonstrating that genes enriched in the macrophage colony stimulating factor pathway (GO-BP) (NES: -3.89, FDR = 0) are mainly downregulated, while in **(C)** are enrichment plots demonstrating that genes enriched in the natural killer cell mediated cytotoxicity pathway (NES: 2.65, FDR < 0.001) are mainly upregulated in ERα KO as compared to control HIC. NES: Normalized enrichment score.

Guided by our results, the ligand-receptor interactions of interest were investigated using BulkSignalR. This analysis identified the CCL5/CCR5 interaction, as the most active (based on statistical analysis) in the “chemokine receptors bind chemokines” pathway (R-HSA-380108) (FDR < 0.01)) (**Supplementary Figure 5A**), consistent with the role this interaction plays in T and NK recruitment and activation (13). The analysis also suggested an IL6ST/LIF interaction in the interleukin 6 family of signaling pathways (FDR < 0.05) **(Supplementary Figure 5B),** and CSF1 interaction with its receptor CSF1R in the myeloid cells pathway (P < 0.05) **(Supplementary Figure 5C)**. Of interest, the data deconvolution platform CIBERSORTx (20) did not detect major differences in the HIC populations within the liver TME in ERα KO mice as compared to the control, with the exception of T follicular helper cells (Tfh) that increased and infiltrating naïve B cells and plasma cells (but not memory B cells) that showed a trend toward increased presence in among HIC cells **(Figure 6).** Of interest, Tfh cells are specialized CD4^+^ Th cells that promote B cell proliferation and maturation and high-affinity antibody production (21). The increase in the fraction of Tfh cells and the trends to increased fractions of naive B and plasma cells (p>0.05<0.01) in the HIC of ERα KO mice are in line with this role.

**Figure 6.**
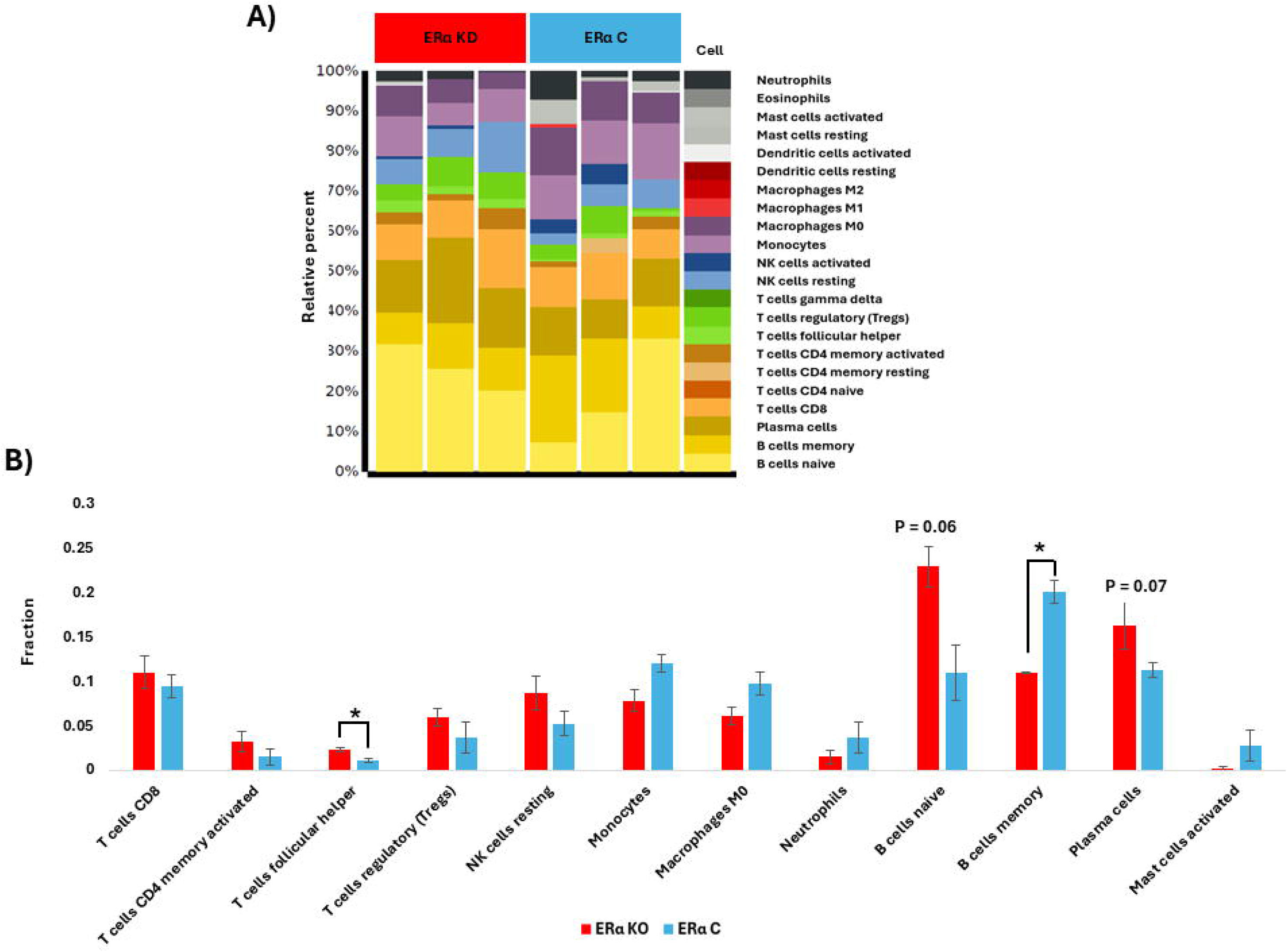
A Cibersortx analysis identifies differences in immune cell subtypes in the TME of liver metastases in ERa KO mice. Shown in (**A**) is a stacked chart depicting the proportions of immune cell subsets predicted from the transcriptomic data for each sample in the comparison groups and in (**B**) the fraction within the total HIC population represented by specific immune cell subsets expressed as means±SEM (n=3). *P < 0.05

## Discussion

We previously reported that estrogen deprivation achieved through ovariectomy or the use of estrogen receptor inhibitors altered the microenvironment of the liver and reduced liver metastases from different tumor types. Because estrogen receptors – ERα in particular, are ubiquitously expressed, we sought here to elucidate the specific contribution of ERs expressed on bone-marrow derived myeloid cells. We show that a selective, inducible deletion of ERα (but not ERβ) on myeloid-derived cells reprogrammed the IME of the liver, resulting in increased T cell mediated cytotoxicity and reduced experimental metastases of colon cancer MC-38 cells. Intriguingly, in mice with a selective deletion of myeloid cell ERβ we did not observed changes in the IME and noted a trend towards increased liver metastases, indicating that the two estrogen receptors played distinct and potentially opposing roles within the TME of the liver.

Our results suggest that the reduction in metastases was linked to a reprogramming of the tumor IME of the liver, whereby the frequency of M0 and M1-like recruited macrophages increased, resulting in an increased M1:M2 macrophage ratio. This indicates that estrogen signaling plays a role in macrophage plasticity and polarization to an anti-inflammatory M2-like phenotype. This is in line with our previously documented findings in ovariectomized mice and also in mice treated with the selective ER degrader Fulvestrant (15). Moreover, the present results are also in line with our findings that in ovariectomized mice, CCL5 levels in the liver significantly increased and that disruption of the CCL5/CCR5 interaction using the CCR5 antagonist Maraviroc reversed the reduction in liver metastases noted in ovariectomized mice (15). Collectively, these data suggest that one of the mechanisms by which ERα can program the IME in the liver is the (down)regulation of the chemokine CCL5 and its interaction with CCR5, as was also predicted by the results of the BulkSignalR analysis.

The source of increased CCL5 production in the livers of ERα KO mice remains to be positively identified. However, our previous results identified local NKT and CD8^+^ T cells as major producers of this chemokine in OVX mice (15). Increased CCL5 production may therefore be linked to increased presence of M1-like macrophages and consequently the increase in the numbers of activated NK and CD8+ T cells.

Among the cytokines downregulated in the HIC of mice with ERα deletion was CSF1 (M-CSF), a well documented transcriptional target of the ERα (22) that is known to play a major role in monocyte/macrophage recruitment and polarization (23–26). This may explain the marked increase observed in the proportion of M0 and M1 among recruited macrophages in the livers of ERα KO mice, as CSF1 is a known inducer of macrophage polarization to the non-inflammatory, M2-like phenotype. Moreover, the decrease in the expressions of other well-known tumor/immunosuppression promoting genes such as *Yap1* (suppressing T-cell function and infiltration) (27) and *Serpine2* (promoting M2 polarization) (28) (**Figure 4D**) are consistent with decreased immunosuppression and increased T cell activation in the ERα KO mice, while the decrease in the expression of Elastase suggests altered neutrophil recruitment to the liver in these mice.

While similarities were found between the effects of ovariectomy and ER blockade on the cytokine/chemokine expression profiles in the liver IME and those observed in mice with myeloid ERα deletions, there were also distinct differences. Thus, while in OVX mice we observed marked increases in the levels of IL-1Ra, CCL2, CXCL9 and CXCL10, mRNA sequencing and/or qPCR did not reveal measurable changes in the expression levels of these mediators in myeloid-ERα KO mice. This suggests that estrogen signaling in other, non-myeloid compartments within the liver ME or resident macrophages such as Kupffer cells contribute to the cytokine/chemokine profile in the TME and thereby to recruitment and activation of immune cells and the cell-cell communications that ultimately shape the microenvironment. This may also explain the more modest reduction in the number of liver metastases in the KO mice, in comparison to the more robust reductions seen in ovariectomized mice or in mice treated with the ER inhibitors tamoxifen and Fulvestrant (14, 15), where a more global blockade of estrogen signaling occurred.

M1 macrophages can mediate T cell recruitment and activation through the release of cytokines (24, 25), among them IL-12 (30) that together with IFNγ, differentiate naive CD4+ T cells into interferon γ producing Th1 cells and enhance the activity of CD8+ T cells (21). Our analysis showed that most transcripts enriched in the IFNγ production-biological process, including *Il12rb2 and Cd40lg* (CD40 Ligand), are upregulated in ERα KO (FDR < 0.05) (**Supplementary Figure 4C**). The increased expressions in ERα KO mice of *Il12rb2* that regulates INFγ production in Th1 T and NK cells and *Cd28* involved in T cell activation (FDR < 0.05) is consistent with our finding of increased activation of these cells in the liver and also suggest heightened responsiveness of these cells to IL-12 (26).

In addition, the increased presence of Tfh cells and a similar trend in naïve B and plasma cells, in ERα KO mice suggests an enhanced humoral immunity in these mice compared to the controls. Taken together, the present findings suggest that ERα expression on the myeloid cells is linked to the plasticity of these cells and their ability to polarize from a proinflammatory to an immunosuppressive, pro-tumor phenotype and that increased M1 macrophage numbers drive an increase in activated T and NK cells. Clinical trials are ongoing with CSF1/CSF1R inhibitors for the treatment of cancer (31). A recent study reported that an immunomodulatory low-dose cyclophosphamide coupled with anti-CSF-1R and anti-PD-1 antibodies was highly effective against aggressive metastatic triple negative breast cancer transplanted in syngeneic mice that present with high macrophage infiltration (32), while single agent targeting of CSF-1R or anti-PD-1 alone were ineffective. Our results suggest that inhibition of ERα signaling may further potentiate the efficacy of treatment with CSF1R/CSF1 inhibitors by blocking macrophage polarization and promoting an anti-tumor immune response.

## Conclusion

This study demonstrated that reduction of ERα in the myeloid cells contributes to reduction in LM and enhance an anti-tumor proinflammatory responses involving M1 macrophage polarization, and recruitment and activation of NK and T cells in the TME. This is mediated, at least in part, through decreased expression of *Csf1* and increased expression of *Ccl5*, in HICs of the TME. Our results suggest that while the reduction in ERα expression in myeloid cells did not alter immune cell recruitment to the liver *per se* it did result in greater anti-tumor activity of the recruited innate and adaptive immune cells, likely due to the altered cytokine/chemokine landscape. In addition, the apparent increase in Tfh cells and their potential effect on antibody producing B cells raise the possibility that in addition to T cell activation B cells also contribute to the reduction in metastases observed in these mice. ER inhibitors may therefore be a useful addition to immunotherapy in the management of liver metastatic disease originating from CRC and potentially other GI cancer, in particular in pre-menopausal female patients.

## Materials and methods

### Mice

All mice experiments were carried out in strict accordance with the guidelines and regulations of the Canadian Council on Animal Care (CCAC) and under the conditions and procedures approved by the Animal Care Committee of McGill University (AUPs- MUHC- 5260). All mice were bred and housed in the animal facility of the RI MUHC. All experiments were performed on 7-16 wk old female mice. The LoxERa/iLysMcre mice were derived by crossing the following two strains both From the Jackson Laboratory: B6 (Cg)-*Esr1^tm4.1Ksk^*/J (JAX stock #032173) and B6.129P2(FVB)-*Lyz2^tm1(cre/ERT2)Grtn^*/J (JAX stock #032291). The ERβ/iLysMcre were obtain by crossing the floxed ERβ (fERβ) mice (a gift from Dr. Teresa A. Milner, Weill Cornell Medical College, NY) (33) and B6.129P2(FVB)-*Lyz2^tm1(cre/ERT2)Grtn^*/J (JAX stock #032291) mice. Genotyping was confirmed using the Kapa HotStart mouse genotyping Kit (KK7352) (Roche), and using the primers recommended by Jackson Laboratory for each strain.

### Cre activation

The TMX induced Cre recombinase activation protocol is illustrated in **Supplementary Figure 1**. TMX (cat#TRC-T00600, Toronto Research Chemicals (TRC)/LGC Standards) was dissolved overnight at 37°C in sterile corn oil (cat# C8267, Sigma) using a shaking incubator (Forma Orbital Shaker, Thermo). The solution was prepared freshly prior to each experiment, stored at 4°C and warmed up immediately before administration by oral gavage (0.1 ml of a 30 mg/ml TMX solution) once daily for 5 days. No adverse effects of this protocol were observed.

### Experimental liver metastasis

Experimental liver metastases were generated by intrasplenic/portal injections of 2X10^5^ MC-38 cells followed by splenectomy, as we previously described (16). Animals were euthanized approximately 3 weeks later and visible metastases on the surface of the liver were enumerated and sized prior to fixation in phosphate buffered formalin. Data from at least two independent experiments were pooled for each group for comparative analyses. Where indicated, the livers were fixed in 10% phosphate buffered formalin for H&E to detect micrometastases and quantify the metastatic burden or perfused with PBS and 4% paraformaldehyde solution (PFA) for immunostaining to identify cell types infiltrating the liver.

### Histology

Livers were placed in 10% phosphate buffered formalin (cat#305510, Fisher Scientific) for 48 hours and then transferred to 70% ethanol and stored at 4°C. Formalin fixed tissue was processed, embedded in paraffin, and cut into 5 µm sections. Hematoxylin and Eosin (H&E) (ThermoFisher Scientific, cat. 7221, 7111) staining was performed according to the clinical laboratory standard.

### Flow cytometry

To analyze early changes in the TIME, mice were injected with 2 × 10^5^ MC-38 cells via the intrasplenic/portal route, and the livers removed 7 or 8 days later. Livers were minced and enzymatically digested for 30 min at 37°C in 0.1% Collagenase IV (from *Clostridium histolyticum*, Sigma-Aldrich) in cRPMI with mixing. The liver homogenates were filtered through a 74 μm mesh filter to remove debris and undigested tissue, the filtrate centrifuged at 300 g in a 4°C centrifuge for 5 minutes and the cells washed twice in cRPMI. Hepatocytes were removed by a 3 min centrifugation at 60 g and the supernatants centrifuged at 300 g for 5 minutes at 4°C and the pellet resuspended in 3 ml ACK solution (ammonium– chloride–potassium) for 2 minutes to remove red blood cells. The cells were then resuspended in PBS and 10^6^ cells immunostained with the indicated antibodies prior to FC. Single cells were gated based on size (forward scatter), granularity (side scatter), and viability using an eFluor 450 fixable dye (eBioscience, ThermoFisher). Data acquisition was with a BD FACS Diva software and the data analyzed using the FlowJo software. All antibodies used in this study are listed in Supplementary Table 1.

### Immunostaining and confocal microscopy

Immunoflorescence staining and confocal imaging were performed as we previously described (15). In brief, the livers were first perfused with 3 ml PBS and then with 3 ml of a 4% PFA (cat#J19943-K2, Thermo Fisher Scientific) solution, placed in 4% PFA for 48 hr and then in 30% sucrose for another 48 hr prior to freezing at -80^0^C until used. For immunostaining, 20 μm cryostat sections were prepared, incubated first in a blocking solution (1% bovine serum albumin and 1% FBS in PBS) and then for 1 h each with the primary antibodies, used at the indicated dilutions (see Supplementary Table 2) followed by the appropriate Alexa Fluor conjugated secondary antibodies (all at room temperature (RT). The sections were mounted in the Prolong Gold antifade reagent (Molecular Probes, Eugene, Oregon, USA) and confocal images were captured with a Zeiss LSM880 laser scanning confocal microscope and analyzed using the QuPath software.

### T cell cytotoxicity assay

To measure cytotoxicity of hepatic CD8^+^ T cells, the mice were injected with 5 × 10^5^ MC-38 cells via the intrasplenic/portal route, and the livers removed 8 days later. The liver homogenates were prepared as per the hepatic lymphocyte isolation protocol (15). Hepatic immune cells were isolated from ERα KO and control mice and CD8+ T cells FACS sorted using the BD FACSAria III. The sorted T cells were then added to 96 well plates for co-incubation with MC-38 cells that were pre-seeded in the plates 24 hours earlier in medium containing 250nM Incucyte® Cytotox Green Dye (Sartorius, 4633) and 30 IU/ml mIL2 (PeproTech, 212-12-5UG). The cell mixture was analyzed continuously in real time using the Incucyte S3 live-cell imaging system with a 4 × objective in basic mode, with an image acquired every 4 hours for each well in both the phase contrast and green fluorescence channels. The green objects per well (dead tumor cells) were recorded at each time point and normalized to time 0 hour.

### Hepatic immune cell isolation and RNAseq

To analyze early transcriptomic changes in the immune TME, mice were injected with 2X10^5^ MC-38 cells via the intrasplenic/portal route, and the livers removed 7 days later. Hepatic immune cells (including lymphocytes and macrophages) from 3 mice in each group (ERα KO, ERβ KO, Cre- controls) were isolated as described in detail previously (15), mixed with Trizol (cat#15596026, ThermoFisher) and stored at -80°C until RNA extraction. Total RNA from the isolated hepatic immune cells was extracted using the Direct-zol RNA MicroPrep (cat#R2060, Zymo Research), as per the manufacturer’s instructions. Next generation RNA sequencing (100 bp in sequence length) was performed by the McGill University and Génome Québec Innovation Centre using the Illumina NovaSeq X Plus platform. The fastq files and transcripts per million (TPM) values for each gene (using Dragen RNA, Illumina) were provided by the Innovation Center. The fastq data were analyzed using the RNAseq pipeline described in detail elsewhere (34, 35). Briefly, quality analysis of the raw data was analyzed with FASTQC (version 11.3) and adapter related sequences were removed using Trim Galore (version 0.6.10) (https://www.bioinformatics.babraham.ac.uk/projects/). The genome sequence of mouse (*Mus musculus*) and its annotation file (Mus_musculus.GRCm39.115) were obtained from ENSEMBL (https://www.ensembl.org/). Reads were aligned to the mouse genome with HISAT2 (version 2.2.1) (36) and read counts were obtained using HTSeq (version 2.0.2) (37), where the intersection-strict mode was applied.

Exploratory analysis was performed with ExpressAnalyst (38). Low variance (15%) and low abundance (n□<□5) counts were filtered. Data were normalized as trimmed mean of M-values and sample distribution was visualized by principal component analysis. Differential gene expression analysis was performed using edgeR (39) and differentially expressed genes (DEGs) with FDR values of less than 0.05 were accepted as statistically significant. Overrepresentation enrichment network and heatmap clustering were then performed on the data. Biological processes and pathways were considered significant when the p value determined by the enrichment analysis was < 0.05. Heatmaps were generated with the pheatmap function in R (version 4.5.0). Estrogen signaling-related genes were obtained from GeneCards (https://www.genecards.org) (40) and those with a relevance score of > 20 that were also identified as DEGs in this study were used for the relevant heatmap. Quantitative PCR was used to confirm altered expression of genes of interest. For gene set enrichment analysis, the software GSEA (version 4.4.0) was used (41). For this analysis, the KEGG database was obtained from a current Bioconductor package, GAGE (https://www.bioconductor.org/packages//2.12/bioc/html/gage.html), and converted to a *gmt file. DEGs (ERα KO vs Control), without FDR cut-off, were ranked by fold change and used in GSEA. Datasets including Hallmark and Gene Ontology (BP) from the Molecular Signatures Database (MSigDB) (https://www.gsea-msigdb.org/gsea/msigdb) were also used. The IDs of the enriched GO (BP) identified by the GSEA analysis (P < 0.05) were extracted from the *json file (provided by MSigDB) and clustered and visualized as a heatmap by the GO_similarity function, using the simplifyEnrichment package (v2.3.0) (42) in R (v4.5.0).

### Cell-to-cell communication analysis

Ligand-receptor cellular networks were inferred using BulkSignalR (v1.2.2) in R (43). The read counts by HTSeq were used as input. The functions findOrthoGenes and convertToHuman were used to obtain an expression matrix to a *H. sapiens* equivalent. The FDR threshold was set to 0.001 to define ligand-receptor interactions and enrichment of Reactome biological pathways (https://reactome.org/).

### Data deconvolution

To predict immune cell populations in the liver TME, the RNAseq data was deconvoluted. To this end, the Ensembl IDs of genes (provided by the sequencing facility) were converted to gene symbols using g:Profiler (44). Genes with transcripts per million (TPM) values were searched against the LM22 matrix (based on 547 genes to distinguish 22 hematopoietic cell phenotypes), using CIBERSORTx with default parameters (20).

### Quantitative PCR (qPCR)

Expression levels of genes of interest, identified by the RNAseq analysis was validated using qPCR. The cDNA was synthesized from extracted RNA using M-MLV Reverse Transcriptase (Invitrogen) and the PCR performed with FastStart Universal SYBR Green Master (Roche) and the primers listed in **Supplementary Table 2,** using the 7500 Real-Time PCR System (Applied Biosystems™). Expression levels in ERα KO mice were compared to ERα C using the 2(-Delta Delta C(T)) method (45) and triplicate samples per group.

### Statistical analysis

The Mann-Whitney test was used to compare metastasis data. The Student’s t test was used for all other comparisons. Statistical analyses and graph generation were performed using R (version 4.5.0).

## Acknowledgments

The authors are indebted to the Histopathology Platform of the RI-MUHC and its manager Ms. Fazila Chouiali for their help with the H&E analysis, to Shibo Feng and Dr. Min Fu of the RI MUHC Molecular Imaging Platform for their technical expertise and help with confocal microscopy, to Hélène Pagé-Veillette, Marie-Helene Lacombe and Ekaterina Iourchenko of the RI MUHC Immunophenotyping Platform for their assistance with flow cytometry and to the McGill University and Génome Québec Innovation Centre for their excellent RNA sequencing service.

## Funding

This work was made possible by grants from the Cancer Research Society, the Canadian Institute of Health Research (CIHR- PJT-169032). O. Haçariz was the recipient of a RI-MUHC postdoctoral fellowship and a Peter Quinlan Fellowship in Oncology, Faculty of Medicine and Health Sciences, McGill University.

### CRediT authorship contribution statement

**Pnina Brodt:** Conceptualization, Writing – review & editing, Supervision, Funding acquisition, Project administration. **Orçun Haçariz:** Writing – review & editing, Funding acquisition, Data curation, Methodology, Validation, Investigation, Formal analysis, Visualization. **Megan Kalaw:** Writing – review & editing, Data curation, Methodology, Validation, Investigation, Formal analysis, Visualization. **Qin Yang:** Writing – review & editing, Data curation, Methodology, Validation, Investigation, Formal analysis, Visualization. **Stephanie Perrino:** Methodology, Data curation.

## Declaration of competing interest

The authors have no competing interests to declare.

## Ethics approval and consent to participate

This study was approved by the RI-MUHC IRB. All tissue acquisition was carried out according to the institutional guidelines. All experimental procedures involving mice were carried out with the approval of the Animal Care committee of McGill University.

## Supplementary Information

Supporting Information is provided in the **Supplementary Information file.**

## Data availability

The authors declare that the data supporting the findings of this study are available within the paper and its Appendix A files. Unprocessed (raw) data can be made available by the authors upon reasonable request. The RNAseq dataset (raw and processed) for this study was deposited in the NCBI Gene Expression Omnibus (GEO) database repository, with the accession number (GSE344047).

## Notes

### Competing Interest Statement

The authors have declared no competing interest.

