## Supplementary Information file for "A conditional, myeloid-cell specific estrogen receptor α deletion reprograms the liver immune microenvironment and impedes the growth of colon carcinoma liver metastases"

**Supplementary Figure 1**

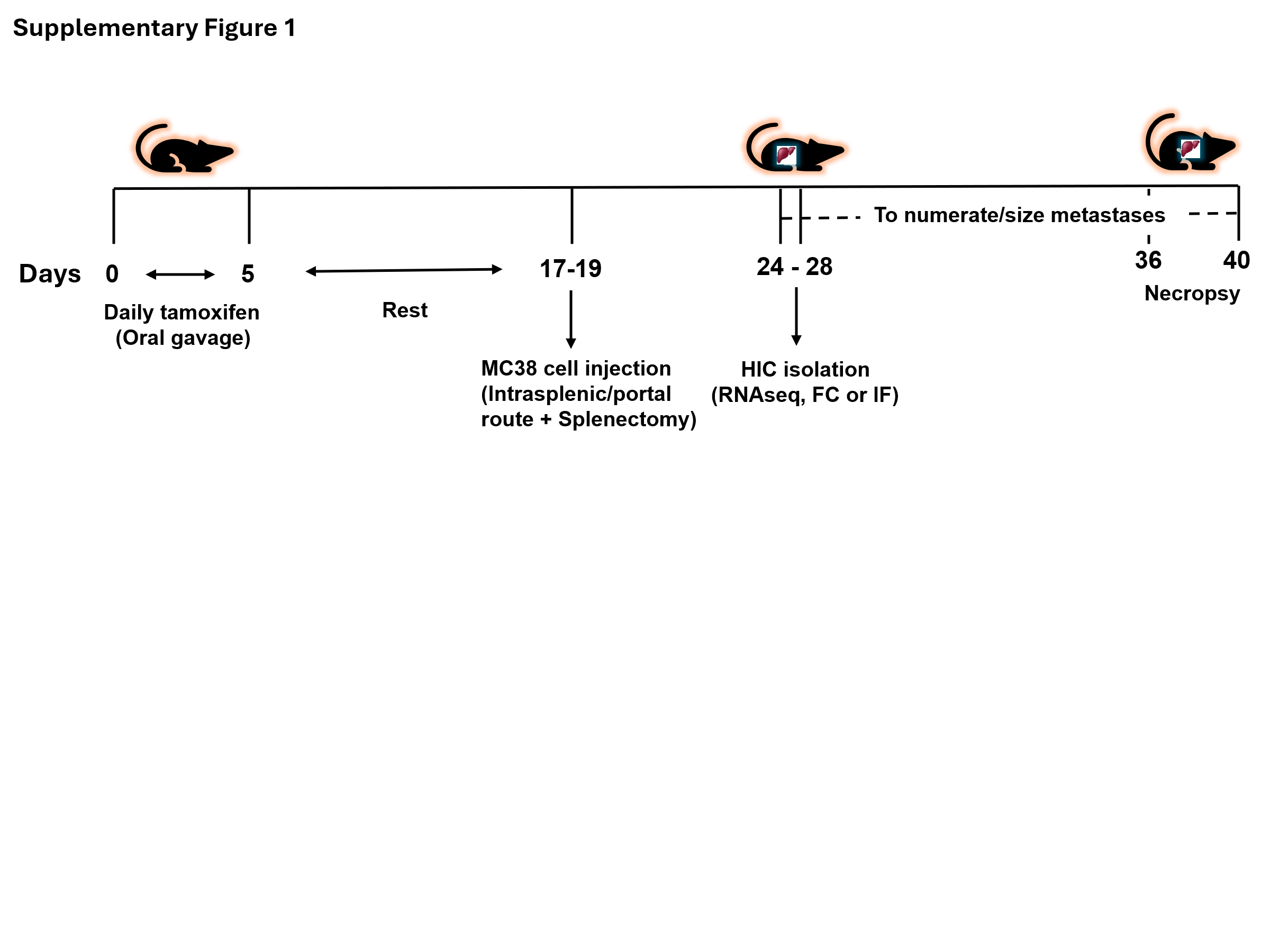

**Supplementary Figure 1. Protocol for Cre recombinase activation and subsequent analyses.** Shown is a schematic representation of the experimental protocol used to activate Cre recombinase and the following experiments. Mice received 0.1 ml of a tamoxifen solution (30 mg/ml) in corn oil by oral gavage, daily for 5 days. Twelve-fourteen days after the last tamoxifen injection (days 17-19), 2 X 10^5^ MC-38 cells (or as indicated) were injected via the intrasplenic/portal route and the spleens removed. Seven-nine days later (or as indicated) the hepatic immune cells were isolated for immune TME analysis (early events) or the mice euthanized 17-23 days later to enumerate and size liver metastases.

**Supplementary Figure 2**

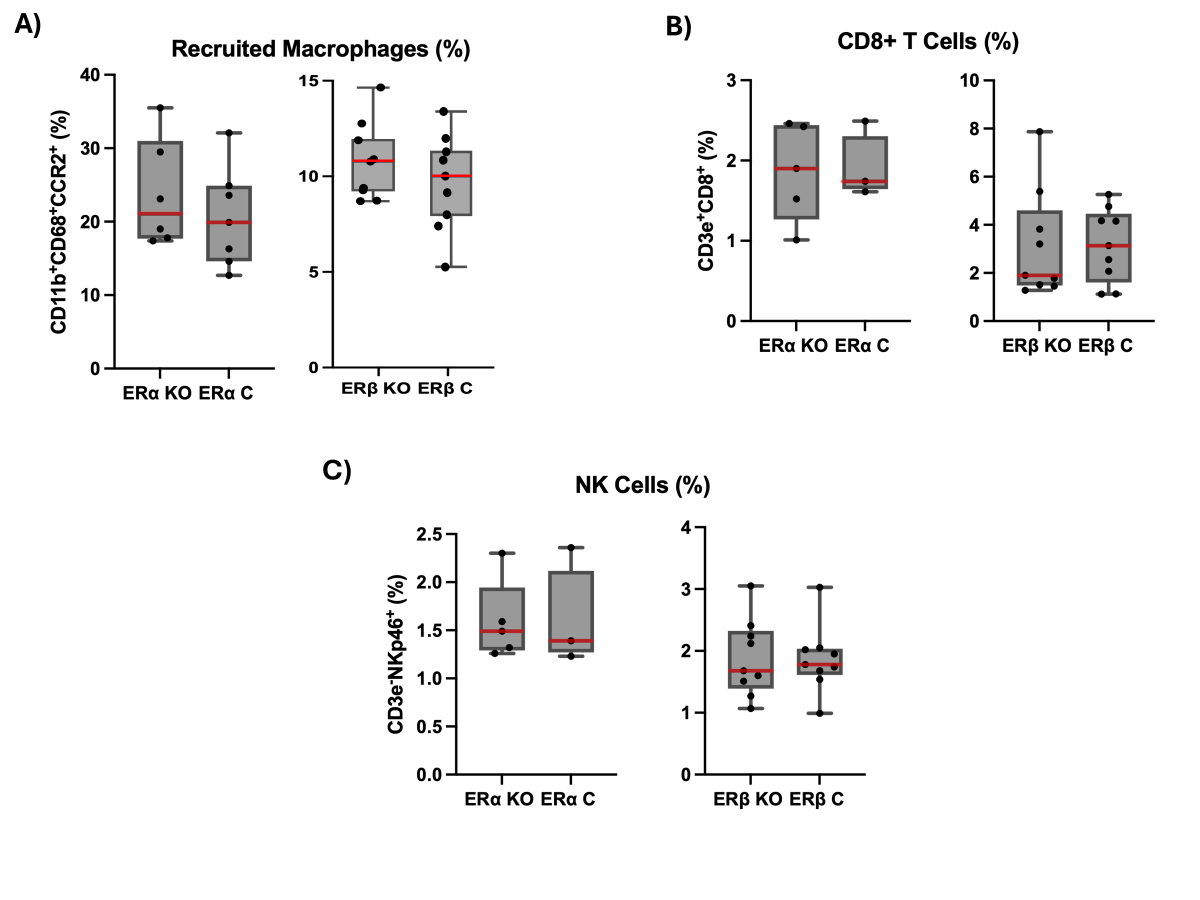

**Supplementary Figure 2. Loss of ERα expression on myeloid cells does not affect their recruitment and the recruitment of lymphoid cells to the liver.** Shown in (**A**) are percent recruited monocyte/macrophages, in (**B)** percent CD3e^+^/CD8b^+^ T cells and in (**C**) percent CD3e^-^/NKp46^+^ NK cells. The results are based on 2 experiments, each with 3-5 mice per group. No significant difference was observed between the groups for any of these immune cell types.

**Supplementary Figure 3**

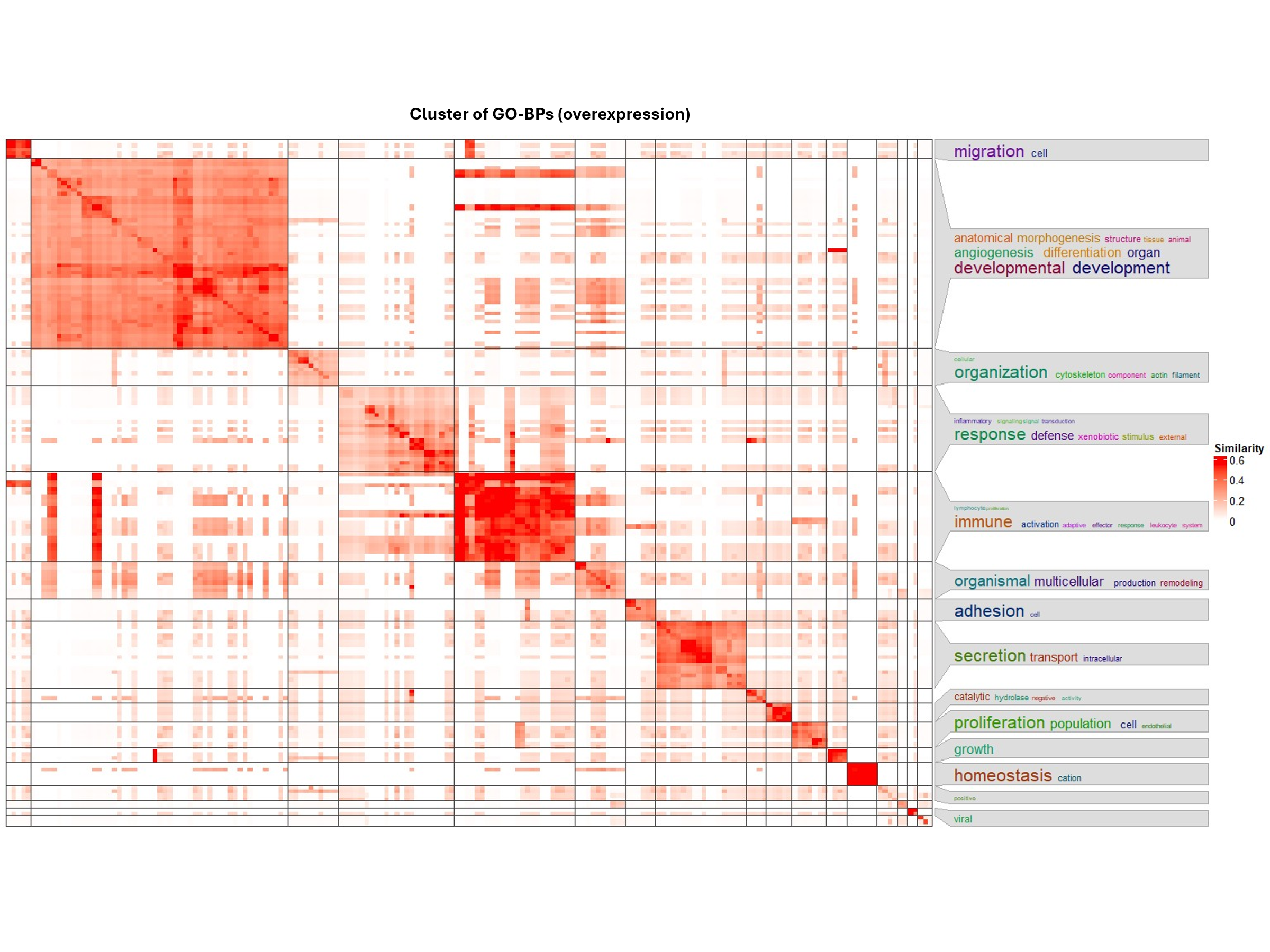

**Supplementary Figure 3. Gene enrichment analysis reveals changes in expression of genes involved in cellular organization and the host immune response.** Shown is a heatmap providing an overview of patterns of biological events in the immune TME, based on clustering of the enriched GO-BPs in ERα KO and the control by simplifyEnrichment (P < 0.05). The biological processes identified appear to be grouped mainly in immune responses, cell migration, development, and secretion.

**Supplementary Figure 4**

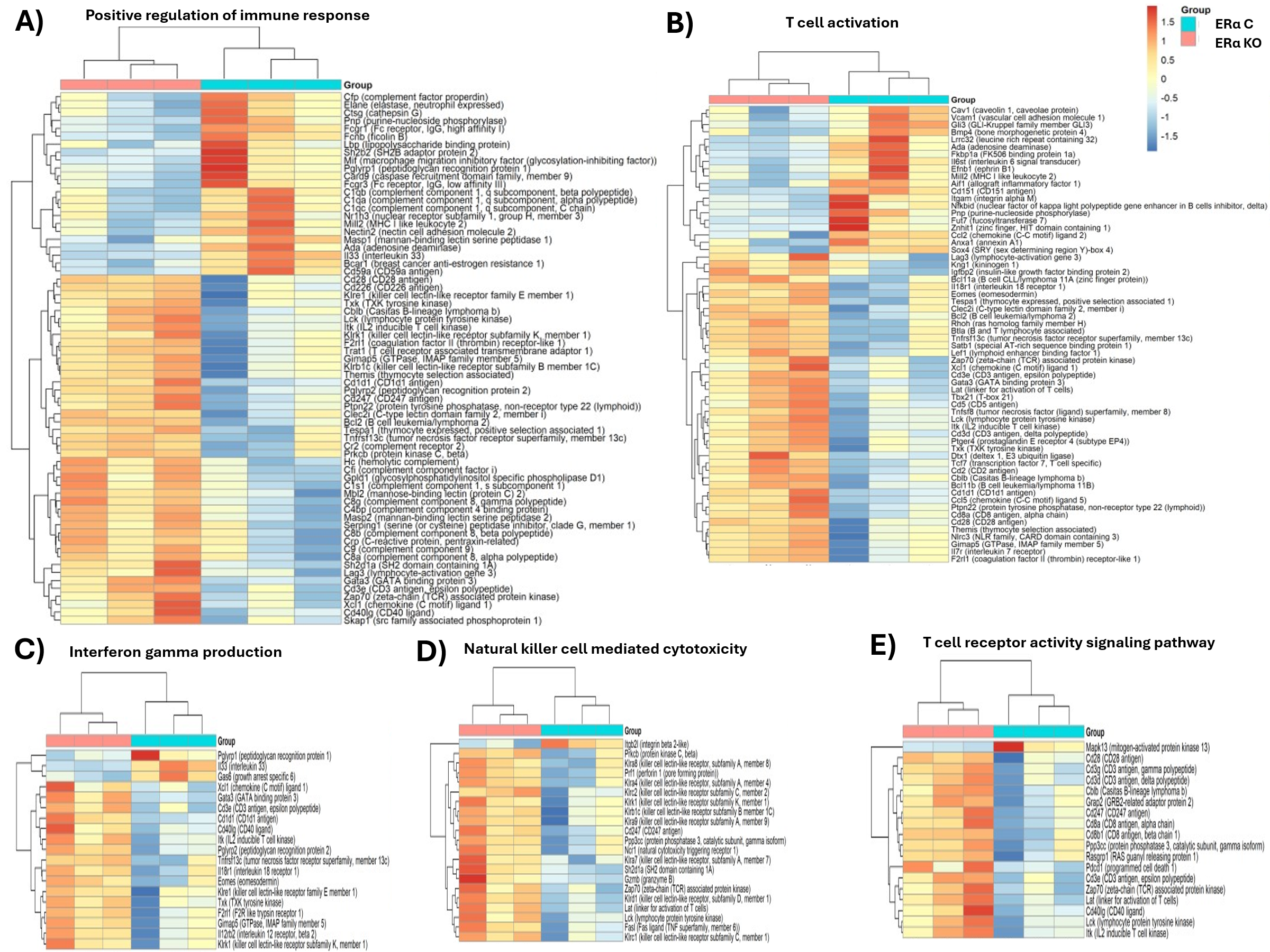
**Supplementary Figure 4. Altered expression of genes mediating anti-tumor immunity in mice with a myeloid-specific ERα deletion.** Shown are heatmaps generated for selected biological processes (GO-BPs) or pathways (KEGG) that are relevant to host anti-tumor immunity. Listed in **(A)** are genes enriched for positive regulaters of the immune response (GO:0050778, FDR < 0.001), in **(B)** T cell activation-related genes (GO:0042110, FDR < 0.01), in **(C)** genes involved in IFNγ production (GO:0032609, FDR < 0.01), in **(D)** genes assocated with NK cell mediated cytotoxicity (mmu04650, FDR < 0.01) and in **(E)** T-cell receptor signaling pathway-associated genes (mmu04660, FDR < 0.05).

**Supplementary Figure 5**

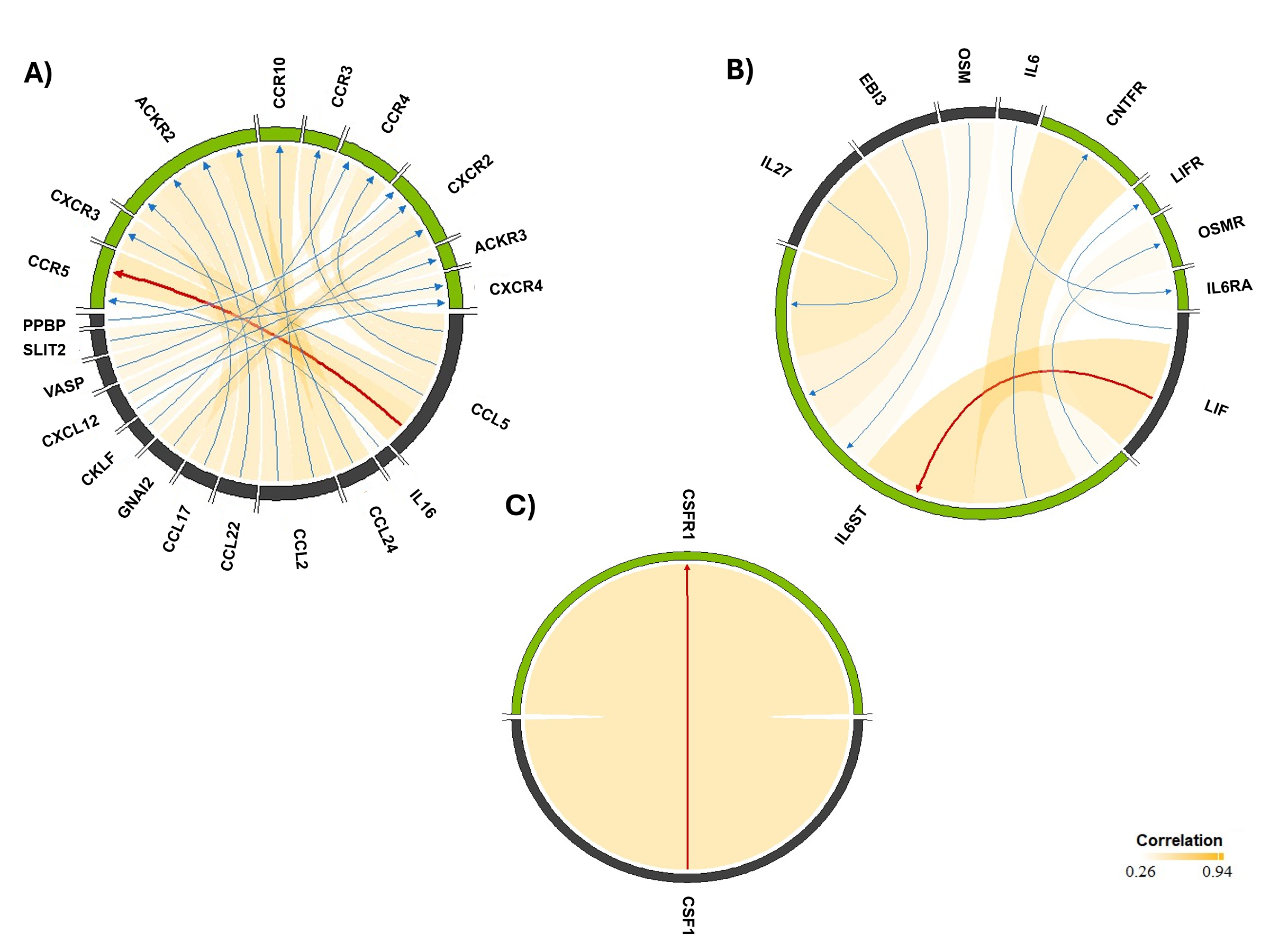

**Supplementary Figure 5. Ligand-receptor interactions in the TME of the liver.** Chord diagrams demonstrate interactions identified by BulkSignalR. Analysis of data shown in **(A)** identifies the interaction between CCL5 and CXCR5 as the most enhanced out of the 17 interactions detected in the [chemokine receptors bind chemokines](https://reactome.org/content/detail/R-HSA-380108) pathways (R-HSA-380108), (FDR < 0.01) in **(B)** IL6ST interaction with LIF, CNTFR and IL27 was the most enhanced of the 8 interactions detected in the interleukin 6 family signaling pathway (R-HSA-6783589) (FDR < 0.01), and in **(C)** CSF1R was identified as the only interactor with CSF1 in the signaling by CSF1 (M-CSF) pathway (P < 0.05). The interactions with the highest correlation scores are highlighted with red arrows.

**Supplementary Table 1**

| **Name** | **Company** | **Cat. Number** |
| --- | --- | --- |
| CD11b-SB702 | eBioscience | 67-0112-82 |
| CD163-SB600 | eBioscience | 15-1631-82 |
| GZB-PE Cy7 | eBioscience | 25-8898-82 |
| NKp46-PE ef610 | eBioscience | 61-3351-82 |
| CD68-AF488 | Biolegend | 137012 |
| CCR2-BV510 | Biolegend | 150617 |
| CD38-PE Cn5 | Abcam | ab25043 |
| CD3e-APC | BD | 553066 |
| CD8b-AF700 | BD | 567737 |
| IFNγ-PE | BD | 554412 |
| CD38 | Proteintech | 60006-1-IG |
| CD68 | Biolegend | 137001 |
| CD163 | Abcam | ab182422 |
| AF568 goat anti-mouse | Invitrogen | A11031 |
| AF488 goat anti-rat | Invitrogen | A11006 |
| AF647 | Invitrogen | A21244 |

**Supplementary Table 1. Antibodies used in this study.** Listed are the antibodies used for flow cytometry all at a dilution of 1:100 and those used for immunofluorescence staining (underlined) all at a dilution of 1:800.

**Supplementary Table 2**

| **Target gene** | **Forward sequence** | **Reverse sequence** |
| --- | --- | --- |
| ***Ccl5*** | GTGCCCACGTCAAGGAGTAT | TTCTCTGGGTTGGCACACAC |
| ***Cxcl10*** | TGGTCTGAGTCCTCGCTCAA | TGATAACCCCTTGGGAAGAT |
| ***Actb*** | AGCCTTCCTTCTTGGGTATGG | GCACTGTGTTGGCATAGAGG |
| ***Ccr5*** | AGCCAGAGGAGGTGAGACAT | GAGCTGAGCCGCAATTTGTT |
| ***Il6st*** | GGGAAAGGAGATGGTTGTGC | GAATTCCGGCCCATCTTTCC |
| ***Csf1*** | GCCTCCTGTTCTACAAGTGGAAG | ACTGGCAGTTCCACCTGTCTGT |

**Supplementary Table 2. Primer sequences.** Listed are the primer sequences used for qPCR analysis.

**Supplementary Data**

**Supplementary Data 1. Differentially expressed genes in ERα KO mice.** Shown is the complete list of DEGs (FDR < 0.05). The number of DEGs in HIC derived from ERα KO (compared to control) was 1368.

**Supplementary Data 2. Enriched datasets based on gene overexpression.** Shown is the complete list of enriched biological processes and pathways identified by GO-BPs and KEGG biological pathways for the indicated comparison groups (P < 0.05). The DEGs with the FDR threshold (<0.05) were subjected to gene enrichment analysis by ExpressAnalyst.

**Supplementary Data 3. Enriched datasets based on GSEA.** Shown is the complete list of positively and negatively enriched biological processes and pathways identified by GO-BP and KEGG in the indicated comparison groups (P < 0.05). All identified DEGs (without FDR cut-off) were ranked by fold change and used by GSEA.
